# A model of coevolution and local adaptation between hosts and parasites in continuous space

**DOI:** 10.1101/2022.06.08.494937

**Authors:** Bob Week, Gideon Bradburd

## Abstract

Previous host-parasite coevolutionary theory has focused on understanding the determinants of local adaptation using spatially discrete models. However, these studies fall short of describing patterns of host-parasite local adaptation across spatial scales. In contrast, empirical work demonstrates patterns of adaptation depend on the scale at which they are measured. Here, we propose a model of host-parasite coevolution in continuous space that naturally leads to a scale-dependent definition of local adaptation and a formal definition for the spatial scale of coevolution. In agreement with empirical findings, our model implies patterns of adaptation vary across spatial scales. When measured on spatial scales shorter than the scale of coevolution, we find the farther dispersing species is locally adapted. However, when measured at longer spatial scales, the opposite pattern is observed. We discuss our results in relation to those found using spatially discrete models and to conclusions drawn from empirical studies, and provide an example of our how our results can be used to inform the design of empirical studies.

## Introduction

Interactions between hosts and parasites have shaped patterns of diversity across all scales of biological organization. For example, coevolution with parasites can alter epidemiological dynamics (Best et al. 2010; Débarre et al. 2012; Lion and Gandon 2015), promote the evolution of sexual reproduction (Otto and Nuismer 2004; Lively 2010), yield novel mutualisms (Yamamura 1993), and influence patterns of speciation across a broad range of taxa (Agrawal and Zhang 2021). In each of these examples, the geography of the interspecific interactions plays a critical role in determining ecological and evolutionary outcomes. In particular, patterns of dispersal in each species have important consequences for the evolution of local adaptation between them (Gandon and Nuismer 2009; Tack et al. 2013). The majority of theoretical studies investigating the determinants of host-parasite local adaptation focus on models in which dispersal occurs between spatially discrete locations. However, most species interactions occur in continuous space, and the dynamics of these spatially continuous systems may be only poorly approximated by discrete space metapopulation models. Therefore, there is a need for models that help us understand, and make predictions for, patterns of host-parasite local adaptation in spatially continuous habitats.

Species that evolve in spatially heterogeneous habitats may exhibit adaptation to local environmental conditions, a phenomenon known as local adaptation. More precisely, local adaptation is the fit between adaptive genetic or phenotypic variation and environmental variation (Kawecki and Ebert 2004). Although the environment is often conceptualized as a set of abiotic factors, coevolving species comprise biotic environmental factors for one another. In addition, the locally adapted species is often considered to be “ahead” in the coevolutionary arms race (Kawecki and Ebert 2004; Greischar and Koskella 2007; Lemoine et al. 2012; Koskella 2014; Pérez-Jvostov et al. 2015; but see Lively 1999; Morran et al. 2014; and Nuismer 2017). Furthermore, the identity of the locally adapted species will depend on the relative spatial scales of genetic or phenotypic variation in each species. In turn, these spatial scales of diversity are determined by the interaction between dispersal and selection (Slatkin 1978). Thus, patterns of host-parasite local adaptation will depend on the relative dispersal abilities and relative strengths of selection in each species. After decades of research on this topic, two general insights into the drivers of host-parasite local adaptation have emerged: (1) the species experiencing stronger selection from the interaction will be locally adapted and (2) for intermediate levels of gene-flow, the species dispersing at a faster rate will be locally adapted (reviewed in Gandon and Nuismer 2009). The explanation for (2) is that intermediate levels of gene-flow promote local genetic diversity without leading to gene-swamping. This increases the rate of adaptation in local populations and gives an edge in the coevolutionary arms race.

Although this result holds under several model variations (Gandon et al. 1996; Gandon 2002; Gandon and Michalakis 2002; Nuismer 2006), there are currently no theoretical results on the role of relative dispersal abilities for local adaptation in continuous space (but see Nuismer et al. 2000; Nuismer et al. 2003 for continuous space models of host-parasite coevolution). Previous models studying the the role of relative dispersal abilities of hosts and parasites have made use of metapopulation models, which lack a notion of geographic distance. Because geographic distance is inherent in continuous space models, we may expect results to depend on the spatial scale at which they are observed. Previous theory of local adaptation of a single species in continuous space suggests that local adaptation increases as the spatial scale of environmental variability increases past that of dispersal (Slatkin 1978; Hadfield 2016). However, because coevolutionary systems are dynamic, it is unclear whether we would expect this result to hold when a coevolving species is treated as the other species’ *environment*. In order to study this question, we require a theoretical model of coevolution in continuous space.

A theoretical model of host-parasite coevolution in continuous space could also provide quantitative predictions for patterns of local adaptation measured at different spatial scales. Several empirical studies have concluded that patterns of adaptation observed in empirical coevolutionary systems depend on the scale at which they are measured (Burdon and Thrall 2000; Tack et al. 2013). For example, Tack et al. (2013) measured parasite adaptation at three spatial scales and found parasite local maladaptation was most apparent at larger spatial scales. We lack a theoretical explanation for why this pattern should emerge, and there is a need to further integrate theoretical predictions and empirical observations of spatial patterns of coevolution. In this paper, we aim to close this gap by analyzing the interaction between host-parasite coevolution, random genetic drift, and gene-flow in a continuous two-dimensional habitat using a quantitative genetic model.

We begin by introducing our model, which is a two-species generalization of Slatkin’s (1978) model of quantitative trait evolution in continuous space. We then outline our analytical approach, which builds on the statistics of spatial autocorrelation (summarized in Box 1) and multivariate Gaussian random fields (summarized in Appendix A). We use this approach to develop a scale-dependent definition of local adaptation in continuous space. Assuming genetic variance and local population densities are constant in space and time, and that coevolution is weak relative to abiotic stabilizing selection, we apply our analytical approach to approximate spatial auto-covariance functions that quantify intraspecific patterns of phenotypic variation. Our approach yields an interspecific spatial cross-covariance function that measures the covariance of host and parasite mean traits as a function of the distance between sampled locations for each species. We propose that the spatial scale of interspecific cross-covariance (described in Box 1) can be understood as the spatial scale of coevolution. Finally, we combine our analytical approach and definition of local adaptation to make predictions for patterns of local adaptation measured at different spatial scales.

### Box 1

What is Spatial Autocorrelation?

Our work is, in part, motivated by the need to understand the effects of spatial autocorrelation on measurements of local adaptation. We therefore provide a brief conceptual explanation of spatial autocorrelation for readers unfamiliar with the concept. Spatial autocorrelation is best summed up by the First Law of Geography: “everything is related to everything else, but near things are more related than distant things” (Tobler 1970). We illustrate a spatially autocorrelated process in Figure 1; note that the colors (denoting values) of nearby locations are more similar than those of more distant locations. The *spatial scale* of autocorrelation describes the geographic distance at which similarity decays, i.e., the geographic distance beyond which values observed at one location are no longer predictive of those observed at another. Figure 2 illustrates spatial patterns with autocorrelation that ranges from weak (top-row) to strong (bottom-row), and over short (top-row) and long (bottom-row) spatial scales.

**Figure 1:**
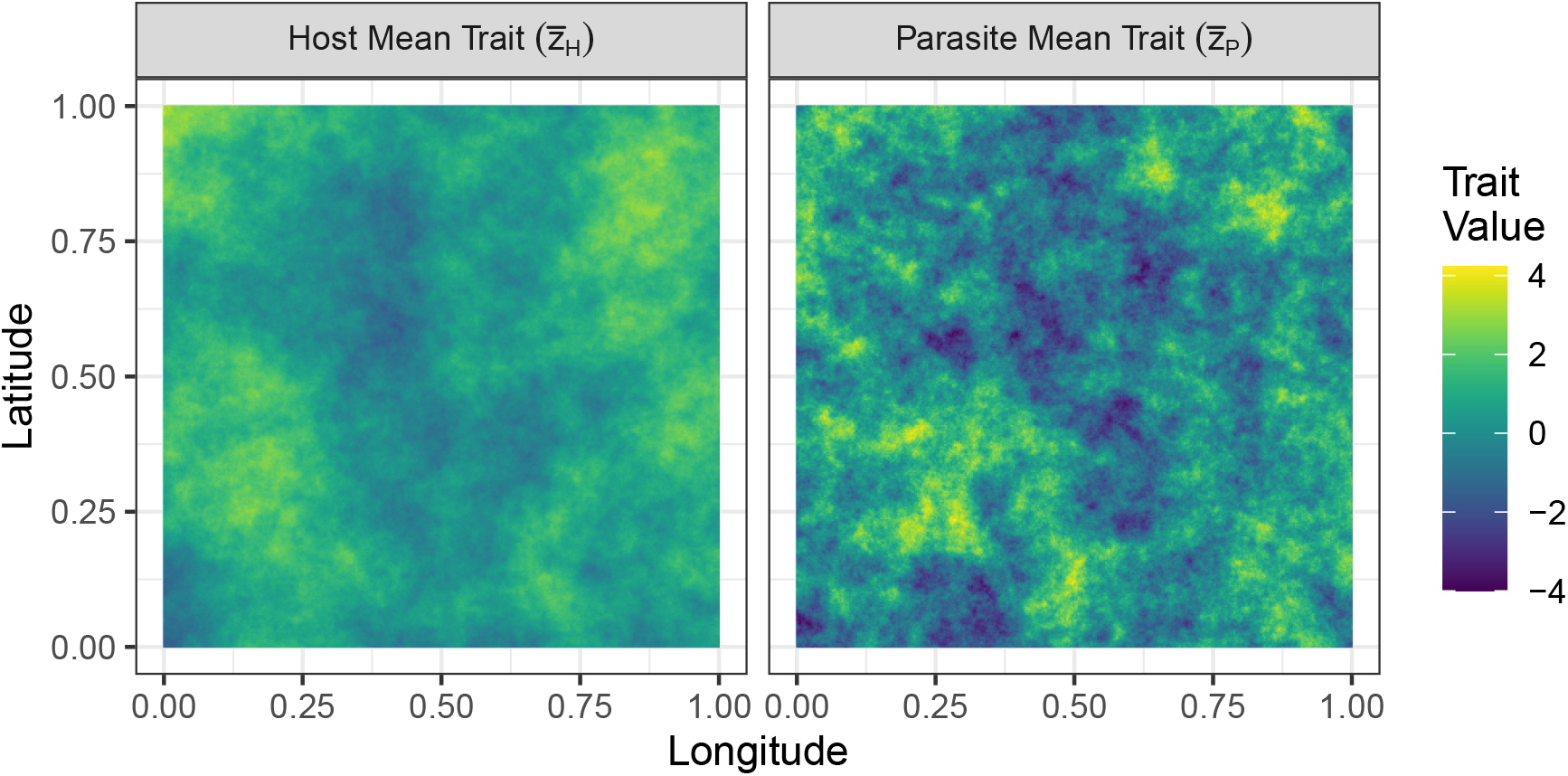
A single realization of our model. Mean trait values for the host (left panel) and parasite (right panel) are plotted as rasterized geographic maps. Here we have set *σ*_*H*_ *> σ*_*P*_. The colocated interspecific correlation of mean trait values 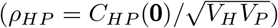 is set to 0.2.

**Figure 2:**
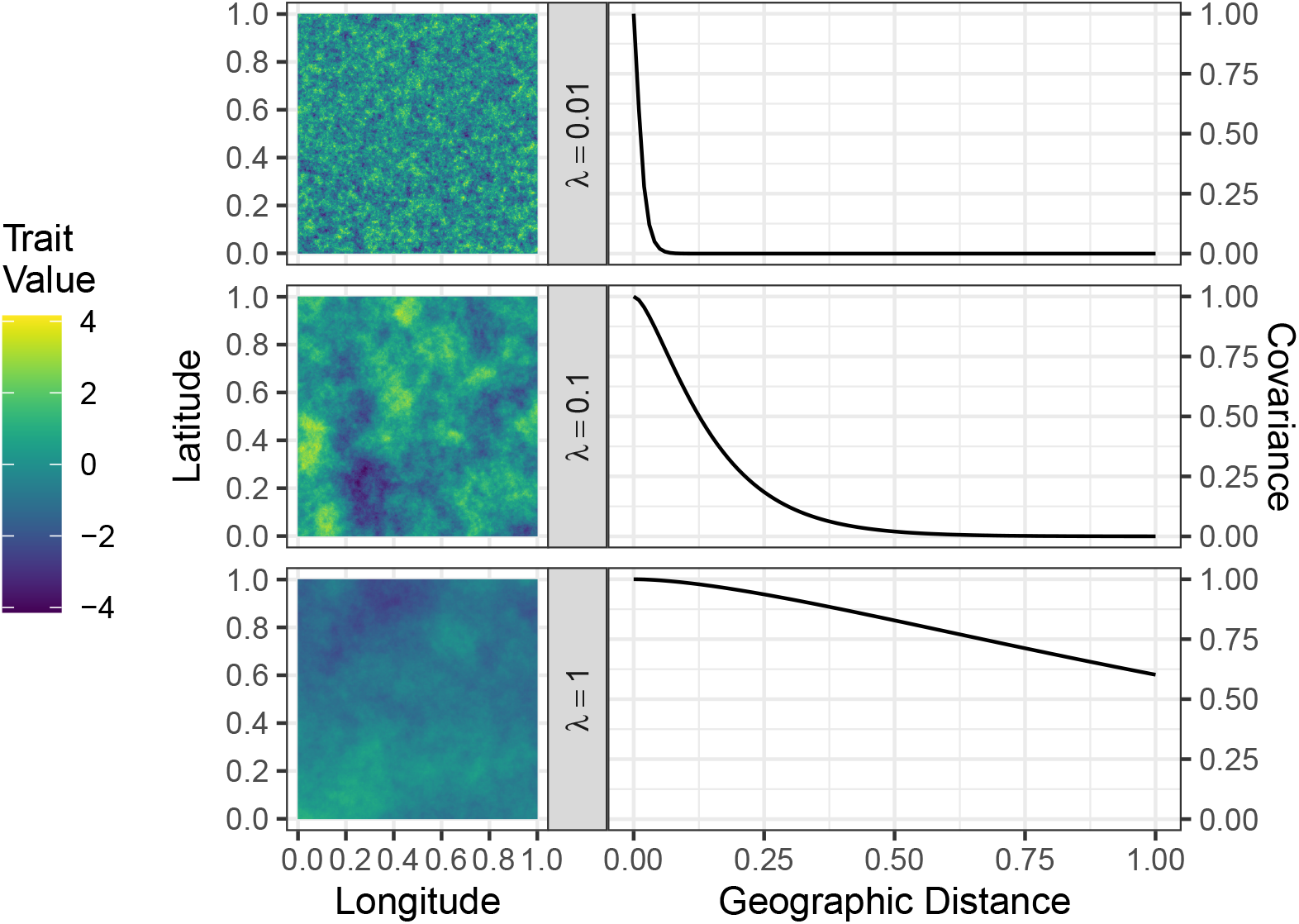
Gaussian random fields and associated covariance functions across three different spatial scales of phenotypic variation *λ* = 0.01, 0.1, 1, with colocated variance *V* = 1.

Throughout this paper, we consider spatial autocorrelation of host and parasite trait values that results from interactions between selection, random genetic drift, and limited dispersal in a continuous geographic landscape. Because coevolutionary patterns involve two spatial patterns (spatial variation in host mean trait and spatial variation in parasite mean trait), we have the additional notion of spatial *cross*-correlation which, loosely speaking, is the autocorrelation *between* the two spatial patterns. For example, if the parasite is locally adapted to the host, parasite trait values will correlate positively with nearby host trait values, but the degree of correlation between host and parasite trait values will decay with the distance between host and parasite populations. In that example, the host and parasite trait values would exhibit positive spatial cross-correlation. Using the framework of random fields, summarized in Appendix A, the magnitude and spatial scale of spatial autocorrelation and spatial cross-correlation can be quantified using spatial covariance and spatial cross-covariance functions (illustrated on the right column of Figure 2).

## Methods

### The Model

Our model tracks the evolution of local mean traits for a pair of species co-distributed across a continuous two-dimensional geographic landscape. For each species, the genomic architecture of their traits is based on an infinitesimal approximation such that the trait of an individual can be thought of as the sum of an infinite number of allelic effects (with no epistasis), each of infinitesimal size (reviewed in Barton et al. 2017). The primary components of our model (selection, reproduction, and dispersal) can be thought of as different stages in the life cycle of an individual. We assume the life cycle begins by determining fitness in response to selective forces, including interspecific interactions, followed by the production of offspring that disperse to new locations, inherit trait values that are normally distributed around parental trait values, and repeat the cycle of life. However, instead of explicitly tracking individuals, our model focuses on the dynamics of mean traits averaged across individuals at each location in geographic space. We assume population densities and genetic variances are constant in space and time; our model is agnostic about whether each species is asexual or sexual.

In this section, we begin by outlining our approach to account for biotic and abiotic selection and to obtain local evolutionary dynamics in response to selection. We then discuss our model of dispersal and random genetic drift before combining these components into our working model. Model parameters are summarized in Table 1.

**Table 1:** Model Parameters & Symbols

| Symbol | Description | Class |
| --- | --- | --- |
| $\bar{z}_H(\mathbf{x})$ | Host Mean Trait at Location $\mathbf{x} \in \mathbb{R}^2$ | Variable |
| $\bar{z}_P(\mathbf{x})$ | Parasite Mean Trait at Location $\mathbf{x} \in \mathbb{R}^2$ | Variable |
| $v_H$ | Local Host Phenotypic Variance | Parameter |
| $v_P$ | Local Parasite Phenotypic Variance | Parameter |
| $G_H$ | Local Host Additive Genetic Variance | Parameter |
| $G_P$ | Local Parasite Additive Genetic Variance | Parameter |
| $N_H$ | Local Host Population Density | Parameter |
| $N_P$ | Local Parasite Population Density | Parameter |
| $\theta_H$ | Host Abiotic Optimal Phenotype | Parameter |
| $\theta_P$ | Parasite Abiotic Optimal Phenotype | Parameter |
| $A_H$ | Host Abiotic Stabilizing Selection | Parameter |
| $A_P$ | Parasite Abiotic Stabilizing Selection | Parameter |
| $B_H$ | Host Biotic Selection | Parameter |
| $B_P$ | Parasite Biotic Selection | Parameter |
| $\sigma_H$ | Host Dispersal Distance | Parameter |
| $\sigma_P$ | Parasite Dispersal Distance | Parameter |
| $\nabla^2$ | Dispersal Operator | Mathematical Operation |
| $\dot{W}_H(\mathbf{x})$ | Space-Time White Noise due to Host Genetic Drift | Input/Forcing |
| $\dot{W}_P(\mathbf{x})$ | Space-Time White Noise due to Parasite Genetic Drift | Input/Forcing |

### Selection

We assume fitness consequences for interactions between hosts and parasites are mediated by the difference in quantitative traits *z*_*H*_ −*z*_*P*_, where *zS* is the trait value of an individual in species *S* = *H, P* for *h*ost and *p*arasite, respectively, such that the probability of infection increases with increasingly similar trait values. This model of biotic selection has been referred to as the trait matching-mismatching model because it leads to evolution of the parasite to match the host trait, while the host evolves to mismatch the parasite trait. We assume the probability of infection is determined by the Gaussian function

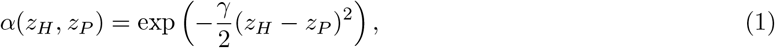

where *γ* ≥ 0 captures the sensitivity of infection probability to trait differences. Given a successful infection, we assume the host incurs a fitness cost *s*_*H*_ ≥0 and the parasite receives a fitness benefit *s*_*P*_ ≥ 0. In addition to biotic selection, we also account for abiotic selection that is stabilizing around an optimum *θ*_*S*_ ∈ ℝ with a strength *A*_*S*_ ≥ 0 for species *S* = *H, P*. Without abiotic stabilizing selection, there is no evolutionary force preventing the host trait from diverging towards infinite values. Because our analysis is performed at equilibrium, we require the host to experience abiotic stabilizing selection to ensure the existence of an equilibrium. Additionally, our approach to computing spatial covariance functions requires that biotic selection be weak relative to abiotic stabilizing selection for both species (see section *Spatial Covariance Functions* below).

Using a series of assumptions, we combine these components of selection to obtain population growth rates, which are then used to obtain expressions for adaptive evolution. First, we assume either that trait values for interacting pairs are sufficiently similar, or that the sensitivity of infection probability to trait differences is sufficiently weak, that *α*(*z*_*H*_, *z*_*P*_) ≈ 1− *γ*(*z*_*H*_ − *z*_*P*_)^2^*/*2. We also assume differences in growth rates caused by selection are small relative to intrinsic growth rates, so that selection is effectively soft (i.e., dynamics of evolution and abundance are approximately decoupled). We assume spatial competition regulates abundance (as in Bolker and Pacala 1997), such that host and parasite population densities *N*_*H*_, *N*_*P*_ are approximately constant in space and time. In addition, we assume parasites encounter local host individuals at random.

Following classical quantitative genetics, we also assume trait distributions of local populations are normal, with mean 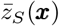, variance *v*_*S*_, and additive genetic variance *G*_*S*_ for species *S* at location *x* ∈ ℝ^2^. In general, this assumption may be violated, as dispersal can lead to skewed trait distributions (Débarre et al. 2015). In Appendix C, we provide an informal argument that our model assumptions imply local trait distributions can be well approximated by normal distributions (and therefore free of skew). To summarize our argument, we find the local trait distributions are exactly normal in the absence of random genetic drift. Our assumptions of weak selection and large local abundance then preserve normality in the stochastic case by preventing substantial eco-evolutionary feedbacks (such as those studied in epidemiological models) and ensuring phenotypic diversity occurs at significantly larger spatial scales than dispersal (so migration occurs between locations with similar trait distributions in each species). As a corollary to this argument, we also assume that phenotypic and additive genetic variances are constant in space and time. Combining these assumptions, we obtain the following growth rates:

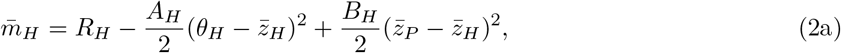

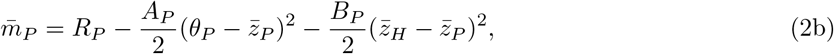

where 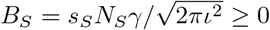 is the strength of biotic selection (with *ι* ≪1 the interaction distance, see Appendix B) and *R*_*S*_ accounts for the intrinsic growth rate (the growth rate when *A*_*S*_ = *B*_*S*_ = 0) and the effects of local trait variance on population growth rates. Further details on model assumptions and calculations made in obtaining these growth rates are provided in Appendix B.

In general, spatially heterogeneous population densities can affect the action of selection (Kirkpatrick and Barton 1997). However, because we assume population densities are spatially homogeneous, we follow Week et al. (2021) to obtain expressions for the local dynamics of mean traits in response to selection. We use 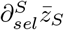 to denote the instantaneous rate of change of 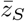in response to selection. Then, because selection in our model is frequency-independent (in the sense that 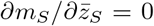), we have 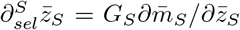. Applying the growth rates obtained above provides

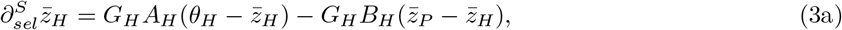

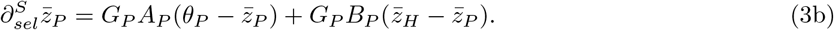

### Drift

Random genetic drift can lead to variation in evolutionary trajectories across spatial locations. In our model, drift provides the ultimate source of geographic variation in phenotypes; this geographic variation then interacts with gene-flow and selection to yield distinct spatial patterns. The classic model for the response of a quantitative character to random genetic drift is given by Lande (1976). This model states that the change in mean trait in response to drift between consecutive, non-overlapping generations follows a normal distribution with variance equal to the ratio of additive genetic variance to effective population size. The continuous time analog of this model is trait evolution following Brownian motion, which has been widely applied as a phenomenological model in the field of phylogenetic comparative methods (Felsenstein 1973; Manceau et al. 2016). Mechanistically, this result has been formalized in continuous time by Week et al. (2021). Denoting 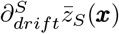 the instantaneous rate of change of the mean trait 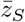 at location *x* in response to drift, Week et al. (2021) found

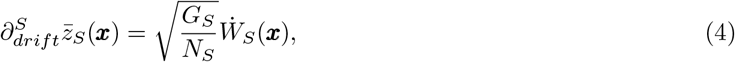

where *G*_*S*_ is the additive genetic variance, *N*_*S*_ is the local population density, and 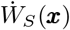 can be thought of informally as an infinitesimal amount of Gaussian noise drawn independently at each spatial location ***x*** ∈ ℝ^2^. Formally, 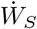 is a space-time white noise process that, when integrated across regions of space and intervals of time, returns a normally distributed variable with mean zero and variance equal to the area of the space-time region integrated over (Walsh 1986).

### Dispersal

We assume displacement between parental and offspring birthplaces follows a bivariate Gaussian distribution centered on zero with latitudinal and longitudinal components drawn independently. These assumptions prevent any net directionality in dispersal. For species *S*, we assume the longitudinal and latitudinal components are drawn with a standard deviation *σ*_*S*_, which we refer to as the dispersal distance (although the expected distance of dispersal is 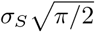).

Following this model, the movement of a single lineage over large spatio-temporal scales will appear as a Brownian motion with rate *σ*_*S*_. Hence, *σ*_*S*_ can be considered both a dispersal distance and a dispersal rate. This aids in comparing our results to previous models of host-parasite local adaptation in discrete space, which consider the rate of dispersal for one species relative to the rate of the other.

Gaussian dispersal leads to change in 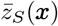 towards a local average, where the scale of “local” is determined by *σ*_*S*_. The local mean trait will increase or decrease depending on the concavity of the spatial mean trait surface. Because the concavity of a surface is quantified by a second spatial derivative, the effect of Gaussian dispersal on the instantaneous rate of change in local mean trait value is related to the second spatial derivative. Denoting 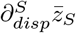the instantaneous rate of change of 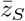 in response to dispersal, we have

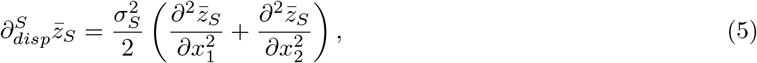

where *x*_1_, *x*_2_ ∈ ℝ are the longitudinal and latitudinal components of ***x*** = (*x*_1_, *x*_2_). For brevity, we often write 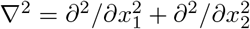so that 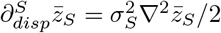. As a technical note, the spatial derivative ∇^2^ must be taken in a weak sense (Evans 2010) because 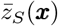 will in general be a non-differentiable function of ***x***.

### Selection, Dispersal, and Drift

We combine the evolutionary forces modeled in the previous sections so that the net rate of change in local mean trait of species *S* is given by 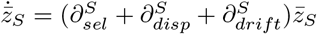. Then, our working model is

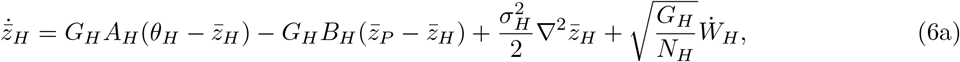

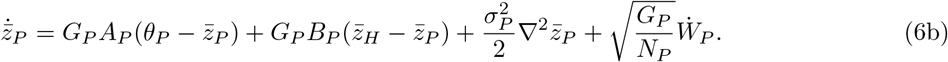

Symbols involved with our model are summarized in Table 1. The system of equations (6) forms a pair of stochastic partial differential equations. In the absence of coevolution, these equations have well-studied Gaussian random fields for equilibrium solutions (Whittle 1954; Whittle 1963; Lindgren et al. 2011). For a summary of this result, see Theorem 7.9 in Lindgren (2012). Because our model of coevolution implies a linear interaction between these fields, the methods used by Whittle (1954; 1963) to study the univariate case can be extended to show that the equilibrium solution of (6) follows an isotropic bivariate Gaussian random field on ℝ^2^. To illustrate our model, a single realization of mean trait values across space for the two species is provided in Figure 1. For the sake of self-containment, we provide a few definitions relevant to Gaussian random fields in Appendix A.

The definition of an isotropic Gaussian random field, summarized in Appendix A, implies equilibrium solutions to our model are completely characterized by five quantities: (1) the expected host mean trait at each location *µ*_*H*_, (2) the expected parasite mean trait at each location *µ*_*P*_, (3) the covariance between host mean traits sampled at any displacement *C*_*H*_ (***x***), (4) the covariance between parasite mean traits sampled at any displacement *C*_*P*_ (***x***), and (5) the cross-covariance between host and parasite mean traits sampled at any displacement *C*_*HP*_ (***x***). Note, the isotropic property of solutions to our model imply that the global variance of mean traits for species *S* (i.e., the variance of an infinitely large sample of mean traits drawn at infinitely large distances from each other) is equal to the colocated variance *V*_*S*_ = *C*_*S*_(0) (which also captures the uncertainty of local trait values due to different possible realizations of drift).

Because our primary interest is in the spatial covariances of mean traits, we ignore the expected values *µ*_*H*_, *µ*_*P*_. In fact, the key aspect of our model that is essential for drawing conclusions on host-parasite local adaptation is the interspecific cross-covariance between traits *C*_*HP*_ (***x***). In the next section, we describe our analytical approach to approximating this cross-covariance function.

### Spatial Covariance Functions

Here we briefly describe our analytical approach to obtaining spatial covariance functions from the system of equations that defines our model, (6). Because our results on local adaptation are obtained from spatial covariance functions, this step in our analysis is fundamental for deriving biological insights from our dynamical model. Essentially, our approach is to compute a spectral representation of our model (i.e., a representation in terms of spatial frequencies), make some simplifications using our assumption that biotic selection is weak relative to abiotic stabilizing selection, then take an inverse transform to obtain the spatial covariance functions. We use the vector ***k*** = (*k*_1_, *k*_2_) to denote spatial frequencies in the two directions (called the wavevector) as opposed to ***x*** = (*x*_1_, *x*_2_), which represents geographic location or displacement. One advantage of this approach is that the spatial derivatives appearing in system (6) become algebraic expressions in terms of spatial frequencies. Another important advantage is the relationship between the spatial covariance function associated with a spatial process and the distribution of harmonic content in that process. The distribution of harmonic content in a spatial process across wavevectors ***k*** is called the power spectrum and can be computed from the frequency space representation (i.e., the Fourier transform) of the process. In turn, the spatial covariance function can then be obtained by taking the inverse Fourier transform of the power spectrum.

The Fourier transform of a function *f* (***x***) can be written 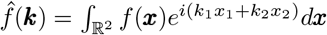. Taking 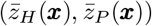 as the equilibrium solution to system (6), we denote the frequency space representation of our model by 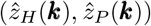. The power spectrum for each species is given by 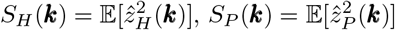, where 𝔼 denotes the expected value across all possible realizations. We can also compute the distribution of harmonic content *between* species, the cross-spectrum, as 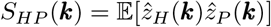. To obtain analytically tractable results, we assume coevolution is weak relative to abiotic stabilizing selection so that *B*_*H*_ ≪*A*_*H*_, *B*_*P*_ ≪*A*_*P*_. This implies that the random mean trait surfaces 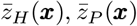 are only weakly coupled. In general, the spatial covariance and spatial cross-covariance functions are given by

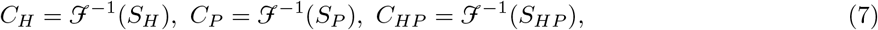

where ℱ^*−*1^ denotes the inverse of the Fourier transform (this is a corollary of Theorems 7.3 and 7.4 in Lindgren 2012). In the Results section, we will see the spatial cross-covariance function *C*_*HP*_ (***x***) plays a central role in our definition of host-parasite local adaptation in continuous space.

### Local Adaptation in Continuous Space

There are two popular definitions of local adaptation at the population level (Kawecki and Ebert 2004; Blanquart et al. 2013; but see Nuismer and Gandon 2008 for a third definition). The first, known as *home vs. away*, is the mean fitness of a population (where mean fitness of a population is defined as the average fitness among all individuals in that population) in its local environment minus the average mean fitness of that population when transplanted to any other location. An alternative definition, known as *local vs. foreign*, is the mean fitness of a population in its local environment minus the average mean fitness of populations transplanted from any other location to that local environment. These definitions are particularly well-suited for metapopulations, comprised of a finite number *K* of discrete locations, because a randomly drawn foreign location (for *home vs. away*) or population (for *local vs. foreign*, but from here on we simply write location) occurs with probability 1*/K*. To directly extend this definition to continuous space, one can consider sampling foreign locations uniformly from a disk in geographic space with radius *r* centered on the focal location, and then take the limit as *r* → ∞. Then one can compute the expected distance between the sampled location and the focal location as a function of *r* and show that this distance diverges towards ∞ as *r* → ∞. Such a definition would therefore lack information about the effects of geographic scale on measurements of local adaptation. Therefore, we introduce a definition that explicitly accounts for the geographic scale at which measurements are taken.

To obtain an index of local adaptation for species distributed continuously in space that accounts for geographic scale, we compute the population growth rate (referred to as a Malthusian growth rate in Crow and Kimura 1970) for a population in its local environment (say at location ***x*** ∈ ℝ^2^) minus the growth rate for the focal population when transplanted to a different location (say ***y*** ∈ ℝ^2^). This definition corresponds to a *home vs. away* definition of local adaptation, as described above. Although classical indices of local adaptation are defined in terms of fitness as expected number of offspring, we chose population growth rate as it leads to relatively simple mathematical expressions. However, given the close correspondence between growth rate and fitness, conclusions drawn using either one should be qualitatively similar.

We denote by *m*_*H*_ (*z*, ***x***) the population growth rate of hosts with trait *z* encountering parasites located at *x*. Similarly, *m*_*P*_ (*z*, ***x***) is the growth rate of parasites with trait *z* encountering hosts located at ***x***. Expressions for these growth rates are given by equations (41) in Appendix B. Setting *φ*_*S*_(*z*, ***x***) as the probability density of trait value *z* in species *S* at location ***x***, the population growth rate for individuals of species *S* transplanted from location ***x*** to location ***y*** is given by

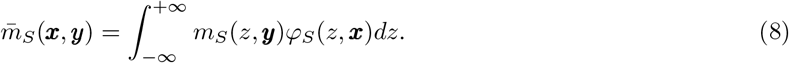

Because our model considers mean traits as random variables, the population growth rates 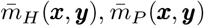 are also random variables. We therefore define local adaptation in terms of expectations of these growth rates. The measure of local adaptation we propose, *ℓ*_*S*_(***x, y***), returns the expected difference between population growth rates for individuals of species *S* drawn from location ***x*** reared locally compared to individuals transplanted to location ***y***. Therefore, this definition explicitly accounts for the spatial distance between locations ***x*** and ***y***. Mathematically, this is expressed as

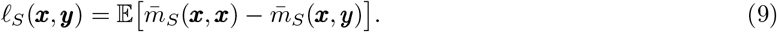

Following this notation, a *local vs. foreign* definition of local adaptation (as described above) would correspond to 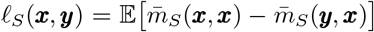. However, under our model, we find 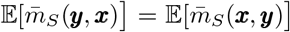 and thus the two definitions coincide. This is a consequence of solutions to our model being spatially isotropic, which is defined in Appendix A.

In the Results section below, we combine this definition of local adaptation with the population growth rates listed in equation (2) to uncover patterns of local adaptation between hosts and parasites coevolving in continuous space.

## Results

### Spatial Covariance Functions

#### Intraspecific Spatial Covariance

Assuming coevolution is weak relative to abiotic stabilizing selection (so that *B*_*H*_ ≪*A*_*H*_ and *B*_*P*_ ≪*A*_*P*_), we obtain simplified expressions for the power spectra. Then taking inverse Fourier transforms of these spectra, we obtain analytic approximations for the (intraspecific) spatial covariance and (interspecific) spatial cross-covariance functions of host and parasite mean trait values. We find spatial covariance functions for the host and parasite respectively take the forms

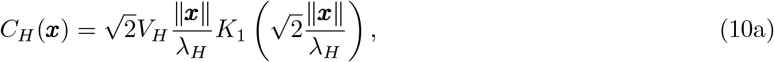

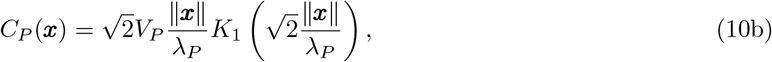

where 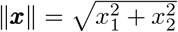is Euclidean distance, *V*_*H*_, *V*_*P*_ are the colocated variances (i.e., *C*_*S*_(0) = *V*_*S*_), *λ*_*H*_, *λ*_*P*_ are the spatial scales of phenotypic isolation-by-distance in each species, and *K*_n_ is the modified Bessel function of the second kind (Abramowitz and Stegun 1965). These spatial covariance functions belong to the class of Matérn covariance functions that have been widely employed in the fields of spatial statistics (Stein 1999; Lindgren et al. 2011) and machine learning (Rasmussen and Williams 2006). In Figure 2, we illustrate the relationship between patterns of phenotypic spatial variation and associated spatial covariance functions by plotting three random fields next to their associated covariance function for three different spatial scales *λ* = 0.01, 0.1, 1 with unit colocated variance *V* = 1.

Our results also demonstrate that the spatial scales of phenotypic isolation-by-distance in the host and parasite can be expressed in terms of model parameters respectively as

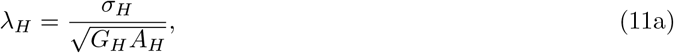

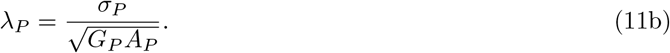

From these expressions, we see that these spatial scales are proportional to the dispersal distances in the respective species; the farther individuals tend to move, the larger the spatial scales one must observe to find significant phenotypic variation. We also see that increased additive genetic variance and abiotic stabilizing selection tend to decrease these spatial scales in each species. Under the assumption of weak abiotic stabilizing selection (which would imply *A* _*H*_, *A*_*P*_ ≪1 and is required in our justification of population growth rates, see Appendix B), our expression for the spatial scale of phenotypic variation coincides with that found by Slatkin (1978).

The colocated phenotypic variances, *V*_*H*_ and *V*_*P*_, represent uncertainty in mean trait value at any particular location due different possible realizations of drift, and should not be confused with the typical notion of phenotypic variance as the variance of trait values among individuals in a population. Because solutions to our model are spatially homogeneous random fields (i.e., they have the same statistical properties at any given spatial location), mean traits of individuals of species *S* sampled at locations separated by distances much greater than *λ*_*S*_ escape the effects of isolation-by-distance and return essentially independent and identically distributed random variables with variances equal to the colocated variance *V*_*S*_. Thus, the colocated variances also provide measures of global diversity of mean traits across space. In terms of our model parameters, the colocated variances can be expressed as

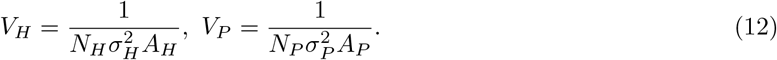

From the expressions for *V*_*H*_ and *V*_*P*_, we see population density *N*_*S*_, dispersal distance *σ*_*S*_, and strength of abiotic stabilizing selection *A*_*S*_ all decrease the overall diversity of mean traits of species *S* across space. Because our model assumes the ultimate source of spatial variation in mean traits is random genetic drift, this explains why increased population density (which decreases the rate of local genetic drift) erodes spatial phenotypic diversity. Similarly, because we assume space extends across the entire plane ℝ^2^, in the limit of infinite dispersal distance (or infinite dispersal rate) we arrive at a panmictic population of infinite size, explaining why increased *σ*_*S*_ decreases *V*_*S*_. These two results can be summarized using Wright’s neighborhood size 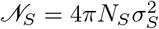, which describes the number of reproducing individuals in species *S* within a disc of radius 2*σ*_*S*_ (the area within which they are effectively panmictic) (Wright 1946; Shirk and Cushman 2014). Because the abiotic optima *θ*_*H*_, *θ*_*P*_ are assumed to be spatially homogeneous, stabilizing selection around these optima erodes spatial variation of mean traits in both species. This explains why *V*_*S*_ is inversely proportional to *A*_*S*_.

### Interspecific Spatial Cross-Covariance

In contrast to the intraspecific spatial covariance functions above, the spatial cross-covariance function, which quantifies interspecific trait covariance measured at two potentially different locations, does not yield a general closed-form solution. Specifically, we find the cross-covariance function is approximated by

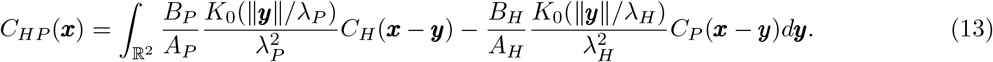

The complexity of this expression creates a challenge for drawing biological conclusions. However, we can gain intuition for this expression by considering two simplifying cases: (1) the case where the host does not disperse and (2) the case where the parasite does not disperse. To obtain these cases, we take the limit *σ*_*S*_ →0, where *S* = *H* for case (1) and *S* = *P* for case (2). For these two cases the cross-covariance function simplifies to

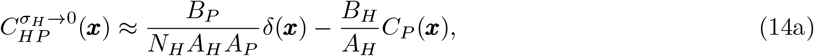

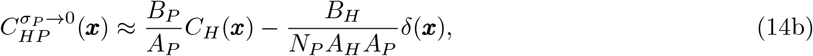

where *δ* is the Dirac delta function (defined such that *δ*(***x***) = **0** for all ***x*** ≠ 0 and 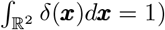 and we have used the superscript *σ*_*S*_ → 0 to label the corresponding cross-covariance function. From these expressions, we see that, in both scenarios, two regimes of spatial covariance emerge. Because one of the species does not disperse, all the effects of coevolution are concentrated at ***x*** = 0. At greater distances, interspecific correlations are maintained by intraspecific autocorrelation in the species that does disperse. These observations suggest the spatial scale of coevolution in each case is zero. However, in general we expect both species to disperse and thus for the spatial scale coevolution to be non-zero. In the next section, we provide a general definition for the spatial scale of coevolution and find that it vanishes whenever the dispersal distance of either species is zero.

### Spatial Scale of Coevolution

Because our cross-covariance function *C*_*HP*_ (***x***) does not suggest a general candidate for the spatial scale of coevolution, we take an approach based on the cross-spectrum *S*_*HP*_ (***k*)**. Specifically, we apply a notion called coherence (Kleiber 2018), which is defined as the linear relationship between two fields at each wavevector ***k***. The coherence function between hosts and parasites in our model is given by

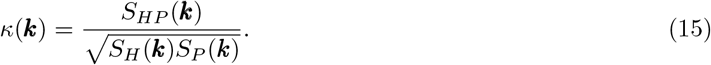

Similar to a correlation coefficient, we have |*κ*(***k***) | ≤1 for all ***k*** ∈ ℝ^2^, where |*κ*(***k***)| denotes absolute value (or modulus when *κ* is a complex number). Building on this notion, we can consider the wavevector of maximum absolute coherence, ***k***_*HP*_ = arg max_***k***_ |*κ* (***k***)|, which is the wavevector at which the two fields are most linearly related. The magnitude 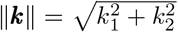 is inversely proportional to a geographic distance. We therefore propose 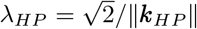as a formal measure for the spatial scale of coevolution, where 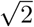 appears as a normalizing constant. Applying this definition to our model, we obtain

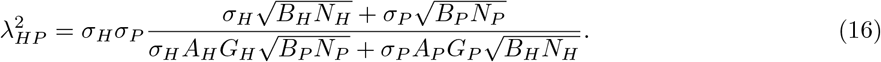

The two special cases treated above can be derived from this general definition. Additionally, if we make all parameters except dispersal distances equal between the two species, we obtain

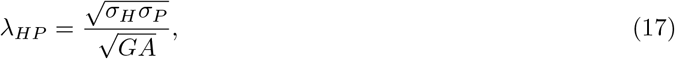

which, in this case, is the geometric mean of *λ*_*H*_ and *λ*_*P*_ (see equations (11a) and (11b)).

If we measure host trait at location ***x*** and parasite trait at location ***y***, this spatial scale of coevolution can be thought of as the distance between ***x*** and ***y*** for which trait values remain statistically dependent. This suggests that *λ*_*HP*_ can be used to formalize the radius of a sampling location where trait values for both species should be measured, i.e., the spatial scale within which phenotypic coevolution will be detectable. Additionally, *λ*_*HP*_ determines the minimum distance between sampling locations above which we obtain approximately statistically independent samples of mean trait pairs. In Figure 3, we illustrate the notions of *λ*_*HP*_ as a radius of a sampling location and as a distance between sampling locations.

**Figure 3:**
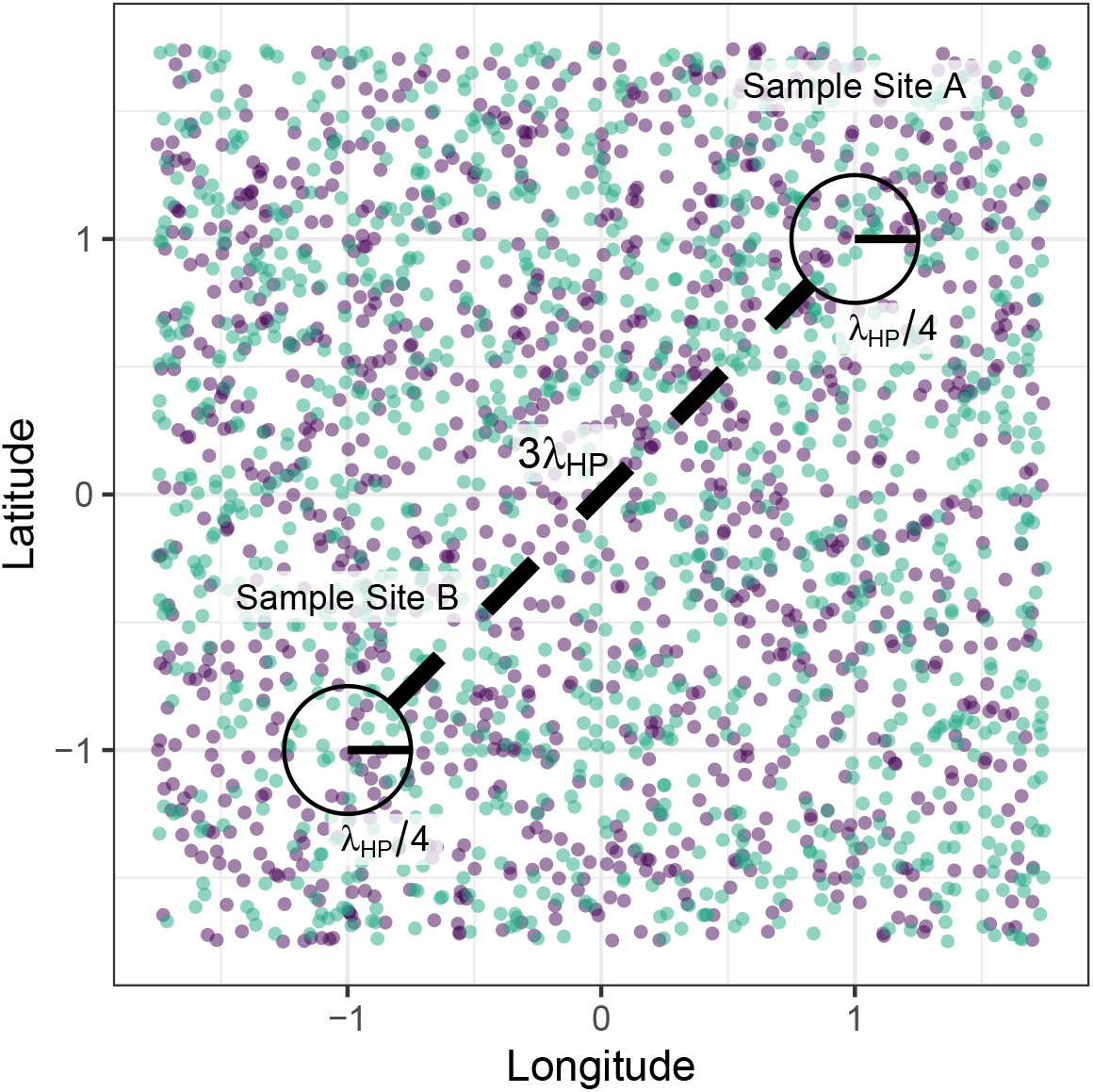
An illustration of the spatial scale of coevolution used to define the radius of sample sites and distance between sample sites. Here, for the sake of illustration, we have arbitrarily set the radius of both sample sites to be equal to *λ*_*HP*_ */*4 (solid lines within each circle) so that trait values measured for a given species within this radius will be highly correlated. In contrast, we have set the distance between sample sites to 3*λ*_*HP*_ (diagonal dotted line) to ensure statistical independence between sites. Decisions on what multiple of *λ*_*HP*_ to use for each distance will depend on the amount of measurement error and statistical non-independence tolerated. Teal dots correspond to host individuals sampled and purple dots correspond to parasite individuals. Following this approach, we would average across sampled hosts/parasites within a defined location to obtain a sample for the host/parasite mean trait at that location. The colocated covariance *C*_*HP*_ (0), which is proportional to parasite local adaptation measured at long distances, can then be approximated by computing the covariance of mean trait pairs sampled at each location. Because *λ*_*HP*_ is not likely known for any system, we describe a heuristical procedure to estimating *C*_*HP*_ (0) in Box 2.

### Patterns of Local Adaptation

#### Analytical Results

Combining our model with our definition of local adaptation in continuous space (defined above in equation (9)), we find

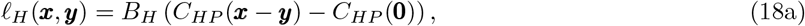

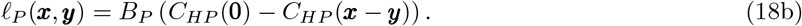

Because all the functions involved with this result depend only on spatial distance *d* ≥ 0, we simplify our notation by writing *ℓ* (*d*) = *ℓ* (***x, y***) when *d* = ||***x −y*** ||. Similarly, we write *C* (*d*), *C* (*d*), and *C* (*d*) for the spatial covariance functions evaluated at a geographic distance *d*. In Figure 4, we present our index of parasite local adaptation as a function of geographic distance across nine combinations of *σ*_*H*_, *σ*_*P*_ = 1, 10, 20 with *N*_*H*_ = *N*_*P*_ = 10, *G*_*H*_ = *G*_*P*_ = 1, *A*_*H*_ = *A*_*P*_ = 0.1, and *B*_*H*_ = *B*_*P*_ = 0.01.

**Figure 4:**
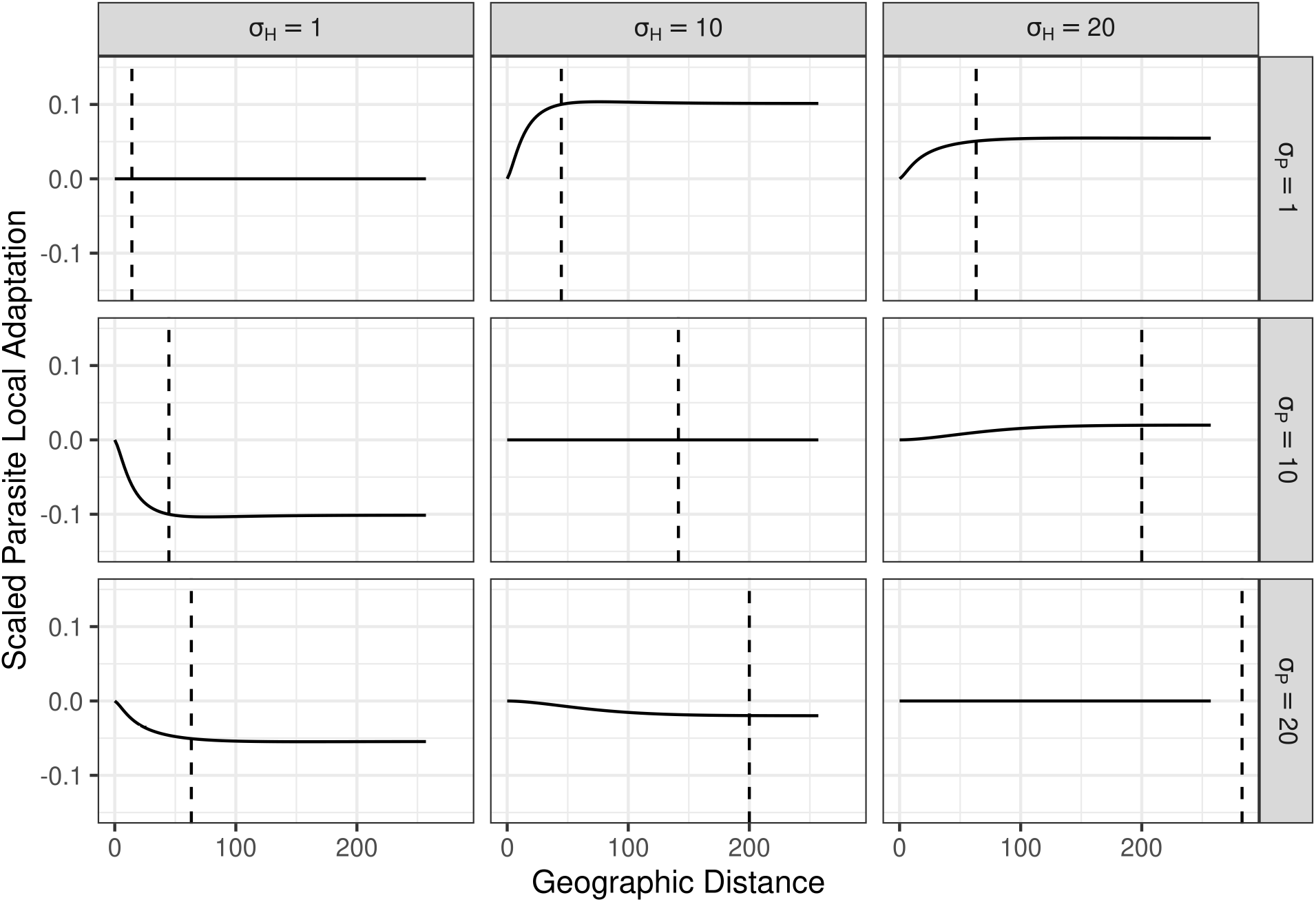
Parasite local adaptation *ℓ*_*P*_ (*d*) as a function of geographic distance, scaled by biotic selection and colocated variances as in equation (19), across nine combinations of *σ*_*H*_ and *σ*_*P*_. Our index of parasite local adaptation under this scaling converges to the colocated interspecific correlation of mean traits asymptotically in *d*. The vertical dashed lines denote the spatial scale of coevolution *λ*_*HP*_ in each case. In the cases where *σ*_*H*_ = *σ*_*P*_ we have *ℓ*_*H*_ (*d*) = *ℓ*_*P*_ (*d*) = 0 for all *d* ≥ 0 so that local adaptation does not emerge at any spatial scale. In the cases where *σ*_*H ≠*_*σ*_*P*_, we see the species with shorter dispersal distance is locally adapted across all distances.

These indices of local adaptation for each species are related by *ℓ*_*P*_ (*d*)*/B*_*P*_ = −*ℓ*_*H*_ (*d*)*/B*_*H*_, which holds when *B*_*P*_, *B*_*H*_ > 0 (i.e., when they coevolve). In addition, because *C*_*HP*_ (*d*) → 0 as *d* → 0, we have lim_*d*→∞_*ℓ*_*P*_(*d*)*/B*_*P*_ = *C*_*HP*_ (0). That is, when parasite local adaptation is measured on large enough distances *d* that *C*_HP_ (*d*) ≈ 0, and divided by the strength of biotic selection *B*_*P*_, it simplifies to the colocated covariance *C*_*HP*_ (0). Furthermore, the colocated interspecific correlation coefficient (defined at the bottom of Appendix A) can be obtained as

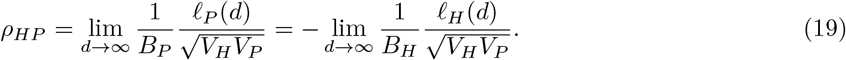

This implies that the species that appears locally adapted when measured at sufficiently large spatial scales is determined by the colocated interspecific spatial correlation of mean traits. Parasite local adaptation at large spatial scales coincides with positive colocated correlations *ρ*_*HP*_ *>* 0 and host local adaptation coincides with negative colocated correlations *ρ*_*HP*_ *<* 0. To understand the role of relative dispersal abilities on the signage of *ρ*_*HP*_, and thus on which species is locally adapted, we evaluated its expression when all model parameters except *σ*_*H*_, *σ*_*P*_ were made equal between the two species. This yields

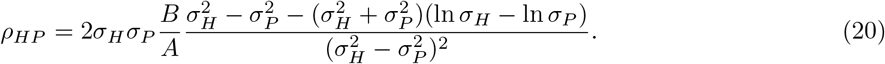

To illustrate this result, in Figure 5 we plotted *ρ*_*HP*_ as a function of *σ*_*P*_ */σ*_*H*_ with all other parameters equal between the two species and *A* = 2*B*. From equation (20) and Figure 5, we see that our model predicts the species with shorter dispersal distance will appear locally adapted at large spatial scales. Additionally, we also see that the magnitude of the colocated interspecific correlation *ρ*_*HP*_, and thus the magnitude of parasite local adaptation, is maximized at intermediate ratios of dispersal distances. In fact, numerical evaluation reveals the formula for *ρ*_*HP*_ given by equation (20) is maximized when *σ*_*P*_ */σ*_*H*_ ≈ 1*/*5 and minimized when *σ*_*P*_ */σ*_*H*_ ≈ 5 for any values of *A* and *B*.

**Figure 5:**
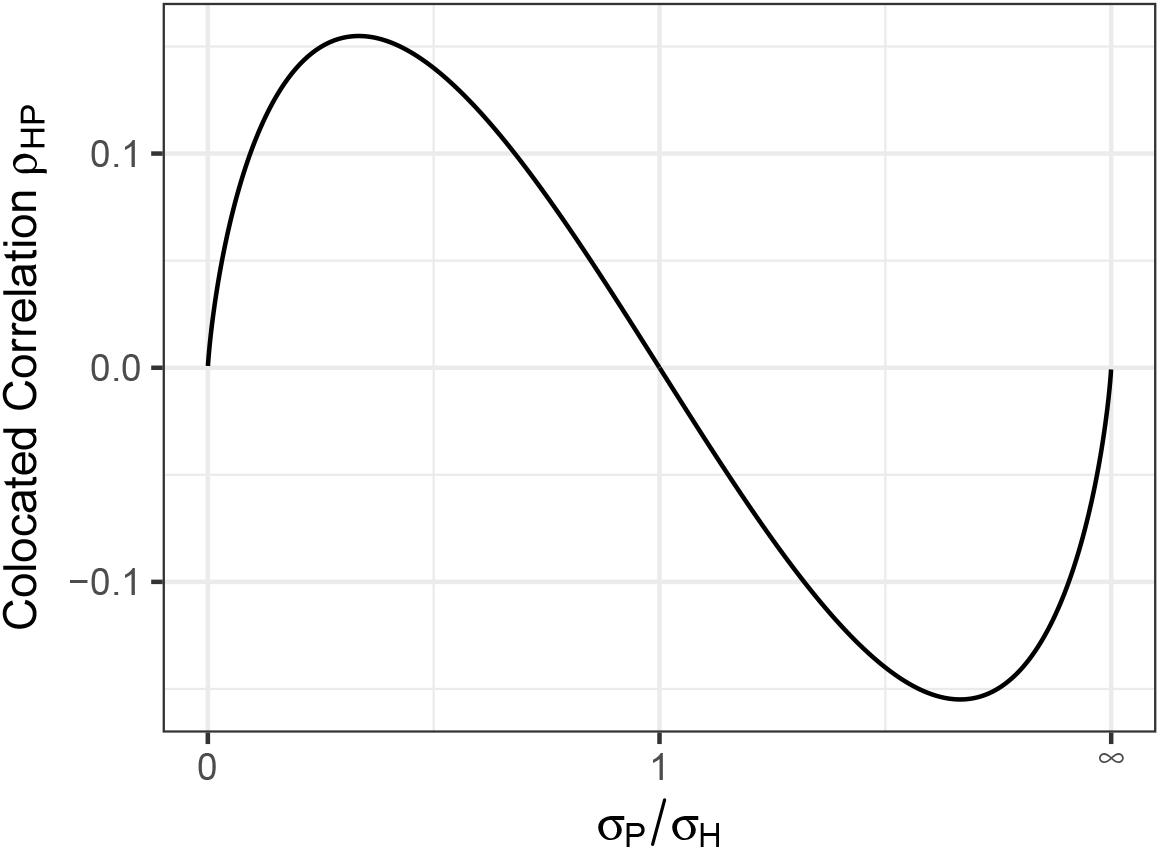
The colocated interspecific spatial correlation coefficient *ρ*_*HP*_ from equation (20), which is proportional to parasite local adaptation, as a function of *σ*_*P*_ */σ*_*H*_ when all other parameters are made equal between the two species and abiotic stabilizing selection is made twice as strong as biotic selection. The *x*-axis has been rescaled to cover the entire range of *σ*_*P*_ */σ*_*H*_.

#### Spatial Scale Dependency

Although the above cases provide important insights by simplifying otherwise complex mathematical expressions, they are limiting in biological scope. For example, in natural systems we might anticipate hosts and parasites to have differing background parameters such as local abundance densities, additive genetic variances, and strengths of abiotic stabilizing selection. Such asymmetries can lead to qualitatively different results compared to our analytical predictions above; in Figure 6 we show two cases that highlight the dependency of the locally adapted species on the spatial scale measured. This result suggests that, to obtain consistent estimates of local adaptation, measurements need to be taken at sufficiently large spatial scales to avoid the effects of spatial autocorrelation.

**Figure 6:**
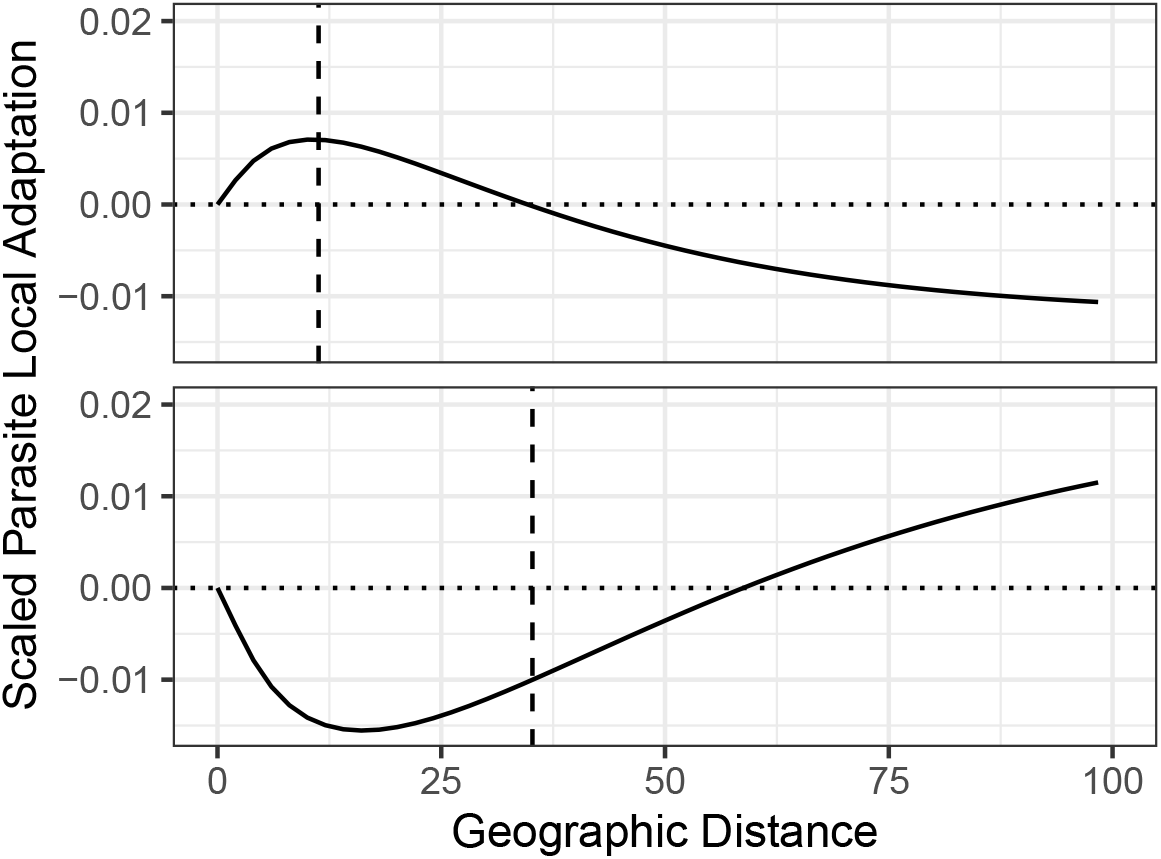
Parasite local adaptation scaled as in Figure 4 and equation (19) for two cases where model parameters are asymmetric between the two species. The two cases presented exemplify the possibility for the identity of the locally adapted species to depend on the spatial scale at which measurements are taken. The vertical dashed lines denote the spatial scale of coevolution, *λ*_*HP*_. For each case we see *λ*_*HP*_ is related to the distance at which the identity of the locally species alternates.

## Discussion

Here, we have extended the study of host-parasite coevolution to populations distributed continuously across a two-dimensional landscape. We investigated the roles of relative dispersal abilities and spatial autocorrelation in determining the identity of the locally adapted species and introduced a formal notion for the spatial scale of coevolution. To conduct this investigation, we developed a model that tracks the evolution of local mean traits of coevolving species in space. This model accounts for host-parasite interactions mediated by a trait-matching/mismatching mechanism, abiotic stabilizing selection, Gaussian dispersal, spatial competition, and random genetic drift. Solutions to this model are Gaussian random fields that are characterized by two intraspecific spatial auto-covariance functions along with an interspecific spatial cross-covariance function. By combining our definition of the spatial scale of coevolution and an index of local adaptation in continuous space with the cross-covariance function obtained from our model, we highlight important considerations for empirical studies of host-parasite coevolution and generate novel insights into the drivers of host-parasite local adaptation.

Although there is already a large body of theory on geographically structured host-parasite coevolution (e.g. Gandon et al. 1996; Nuismer et al. 2000; Gandon 2002; Gandon and Michalakis 2002; Nuismer et al. 2003; Nuismer 2006; Ridenhour and Nuismer 2007; Gandon and Nuismer 2009; Débarre et al. 2012; Lion and Gandon 2015), our model is, to our knowledge, the first to study the interactions of selection, continuous geographic structure, and random genetic drift on the maintenance of phenotypic variation and local adaptation. Previous theory based on a population genetic model has shown parasite local adaptation can be written as the product of a biotic selection parameter and a spatial interspecific covariance of allele frequencies (Nuismer 2017). In analogy to this result, we found parasite local adaptation can be similarly written using a quantitative genetic model, but with mean trait values in place of allele frequencies. However, under our continuous space model, this result holds only when local adaptation is measured across large enough spatial distances to avoid the effects of spatial autocorrelation. As Figure 6 demonstrates, measuring local adaptation on spatial scales at which the effects of autocorrelation remain significant can result in a reversal of the species identified as locally adapted (compared to measurements made at larger spatial scales). This result may explain the spatial scale dependency of parasite adaptation that has been observed in empirically studied systems (e.g. Burdon and Thrall 2000; Tack et al. 2013). However, future work is needed to better understand the conditions under which this phenomenon occurs.

We also found that the relative dispersal abilities of host and parasite can determine which species is locally adapted. Specifically, using our index of local adaptation at large spatial scales, we found that, when all other parameters are equal, the species with the shorter dispersal distance tends to be the one that is locally adapted (see Figure 5). This finding contrasts with previous results that the species with a greater migration rate is the one that is locally adapted (so long as the rate is not so high that it swamps local adaptation, see Gandon and Nuismer 2009). The discrepancy between these results can be explained by one of several differences between our model and previous models. First, our model assumptions imply that genetic variances are constant in space and time, while previous models (of allele frequencies, e.g., Gandon et al. 1996; Nuismer et al. 2000; Gandon 2002; Gandon and Michalakis 2002; Nuismer et al. 2003; Nuismer 2006; Gandon and Nuismer 2009) predict these variances to vary in space and time. Because the evolutionary response to selection is mediated by the amount of genetic variation in a population, changes in genetic variation in time and space can result in qualitatively different evolutionary outcomes compared to when such variation is held constant. Second, it is possible that continuous versus discrete geographic structure explains this discrepancy. Previous quantitative genetic models of local adaptation in a single species to an abiotic environment with continuous spatial structure show local adaptation occurs when the spatial scale of adaptive variation is shorter than the spatial scale of environmental variation (Slatkin 1978; Hadfield 2016). Extending this intuition to a pair of interacting species, we might hypothesize that, all else equal, the species with shorter dispersal distance will tend to be locally adapted. However, to formally show whether our result is a consequence of continuous versus discrete spatial structure or of evolving versus static genetic variance requires further investigation. The extension of our model to freely evolving genetic variances is analytically intractable, so such an investigation would likely be most efficiently conducted using individual-based simulations.

In addition to novel insights into host-parasite local adaptation, our model also suggests a metric for the spatial scale of coevolution. Using a notion called coherence, which quantifies the linear relationship between two random fields across spatial frequencies, we defined the spatial scale of coevolution *λ*_*HP*_ in equation (16) as the geographic distance associated with the spatial frequency of maximum coherence (because distance and frequency are inversely related). This definition is sufficiently general to apply to any form of interspecific interaction, so long as the interacting species are co-distributed across a continuous landscape. Assuming all model parameters are equal for the two species except the dispersal distances, the expression for *λ*_*HP*_ simplifies to the geometric mean of the two intraspecific scales of phenotypic variation 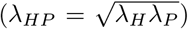. Because the spatial scale of coevolution quantifies the distance at which host and parasite traits are statistically dependent, it is important to consider when measuring local adaptation in empirical studies (see Figure 3). To demonstrate how this information can inform the design of empirical studies, we describe an example in Box 2.

### Box 2

Implication of *λ*_*HP*_ for Empirical Work

Here, we summarize the implications of the spatial scale of coevolution *λ*_*HP*_, defined by equation (16), for the design of empirical studies of coevolution, and provide a concrete example for how we envision the utility of *λ*_*HP*_. Because *λ*_*HP*_ quantifies the distance at which host and parasite traits are correlated, it can be used to determine the radius of sampling areas and distances between sampling locations that together maximize the signal of coevolution in the collected data and minimize spurious signals due to spatial autocorrelation (as illustrated in Figure 3). More specifically, if the spatial scale of coevolution were known, it could be used to inform two aspects of sampling design:

1. The radius of a sampling location (where both host and parasite traits are measured from individuals to estimate local mean trait values) should be made sufficiently short relative to *λ*_*HP*_ to minimize spatial variation that weakens the signal of coevolution. If the radius of a sampling location is too large, individual trait values included in estimates of local mean traits may only be loosely cross-correlated and thereby weaken the signal of coevolution in the data.
2. The distance between locations (across which a correlation of local mean trait pairs can be calculated) should be made sufficiently large relative to *λ*_*HP*_ to minimize spatial autocorrelation of estimated mean trait values. If the distance between a pair of sampling locations is too short, local mean traits estimated may be spatially autocorrelated, which can lead to a reversal of the species identified as locally adapted (see Figure 6).

However, because *λ*_*HP*_ is not likely known for any given host-parasite system, we propose a heuristic approach. Suppose individuals have been sampled uniformly from across their ranges for each species. The question now is how to subset the data into populations such that spatial structure is minimized *within* populations and maximized *between* populations. With geo-referenced trait data for individuals, well-established statistical methods, such as the RFfit function from the R package RandomFields, can be employed to fit intraspecific spatial covariance functions similar to those predicted by our model (the system of equations (10a) above). Because these data are measured from individuals, the spatial covariance functions fitted to the data will slightly differ from those predicted. Regardless, this approach provides a means to obtain rough estimates for the intraspecific spatial scales of phenotypic variation, *λ*_*H*_ and *λ*_*P*_. In turn, because the spatial scale of coevolution *λ*_*HP*_ lies somewhere between the intraspecific spatial scales, *λ*_*H*_ and *λ*_*P*_ provide upper and lower bounds on *λ*_*HP*_. Therefore, the shorter of the two can be used to determine the radius of populations and the longer used to determine the distance between populations. The spatial correlation function for species *S, ρ*_*S*_(*d*), predicted by our model takes the values *ρ*_*S*_ (3*λ*_*S*_) ≈ 0.04 and *ρ*_*S*_ (*λ*_*S*_ */*4) ≈ 0.94. Therefore, populations that are separated by 3*λ*_*S*_ will be approximately independent, taking *λ*_*S*_ as the larger of *λ*_*H*_, *λ*_*P*_. Similarly, taking *λ*_*S*_ as the shorter of *λ*_*H*_, *λ*_*P*_, populations with a radius of *λ*_*S*_*/*4 should have little spatial structure. With these heuristics in place, we can then subset the trait data for individuals into populations (as in Figure 3), compute the mean traits for each species at each population, and, finally, compute a spatial covariance of mean trait pairs across populations that should be approximately free of the effects of spatial autocorrelation. In turn, under our model, this covariance provides a consistent estimator for parasite local adaptation (equation 18b). We provide a short tutorial for this approach at the *GitHub* repository https://github.com/bobweek/interspecific-cov-est.

By combining coevolutionary theory, stochastic partial differential equations, and spatial statistics, we are able to generate new theoretical insights into the mechanics of host-parasite coevolution and local adaptation. However, these results only scratch the surface of possibilities that can be obtained using our novel analytical approach. For example, heavy-tailed dispersal kernels, which are likely common in nature (Houtan et al. 2007; Bullock et al. 2016; García and Borda-de-Água 2016; Jordano 2016), can be incorporated into our model by replacing the dispersal operator ∇^2^ with fractional spatial derivatives (for mathematical details see Laskin 2000; Bayın 2016). Because fractional derivatives are defined in terms of spectral representations, our approach, which recovers covariance functions from spectral representations, is particularly well-suited to this generalization of our model. Another direction this work can be taken is to consider spatial variation of the abiotic environment by modelling the abiotic optima *θ*_*H*_ (***x***), *θ*_*P*_ (***x***) as additional Gaussian random fields (making for four random fields in total). The degree to which the two species experience the same environmental variables can be set by specifying the colocated correlation between *θ*_*H*_ (***x***) and *θ*_*P*_ (***x***). Hu et al. (2013) provides further information on the mathematical details involved with both of these extensions. Lastly, random field models of coevolving species such as ours can be used to extend previous statistical methods developed to measure coevolution from spatially structured phenotypic data (Nuismer and Week 2019; Week and Nuismer 2019). Because these previous methods relied on models that assume discrete spatial structure, they ignore geographic distances between locations. By incorporating random field models, future coevolutionary methods can make use of this information to sharpen inferences on the coevolutionary process.

Our work here provides initial steps towards understanding coevolution and local adaptation in continuous space using an analytically tractable mathematical model. Although our model is relatively simple (in that it ignores selection mosaics and feedbacks with abundance dynamics), it makes testable predictions for spatial patterns of phenotypic diversity and local adaptation resulting from the interplay of coevolution, dispersal, and random genetic drift. Taken together, this work provides a novel analytical approach to discover new theoretical results and deepens our understanding of host-parasite local adaptation.

## Acknowledgments

We thank Anurag Agrawal, Ailene MacPherson, Scott Nuismer, Peter Ralph, and Victoria Caudill for their thoughtful and detailed feedback on a previous draft of this manuscript, which greatly improved this project. Research reported in this publication was supported by the National Institute Of General Medical Sciences of the National Institutes of Health under Award Number R35GM137919 (awarded to G.S.B.). The content is solely the responsibility of the authors and does not necessarily represent the official views of the NIH.

## Appendix

### A Gaussian Random Fields and Spatial Covariance Functions

Because our results are obtained from spatial covariance functions associated with Gaussian random fields (GRFs), we provide some relevant definitions here for the sake of self-containment. Our primary reference is Rue and Held (2005). We begin by defining univariate GRFs before proceeding to multivariate GRFs.

A univariate GRF *F* is completely characterized by its mean *µ*(***x***) = 𝔼 [*F* (***x***)] and spatial covariance *C*(***x, y***) = 𝔼 [(*µ*(***x***) −*F* (***x***))(*µ*(***y***) −*F* (***y***))] functions, where ***x, y*** are geographical locations. For any set of *n* locations ***x***_1_, …, ***x***_*n*_, the *n*-dimensional random vector (*F* (***x***_1_), …, *F* (***x***_*n*_)) has a multivariate normal distribution with mean vector (*µ*(***x***_1_), …, *µ*(***x***_*n*_)) and covariance matrix

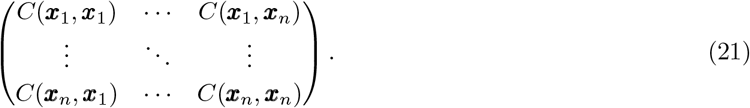

The GRF *F* is called *homogeneous* if its mean is constant across all locations (so that *µ*(***x***_1_) = *µ*(***x***_2_) for any locations ***x***_1_, ***x***_2_) and if its covariance function depends only on the difference between its arguments (so that *C*(***x***_1_, ***y***_1_) = *C*(***x***_2_, ***y***_2_) whenever ***x***_1_ − ***y***_1_ = ***x***_2_ − ***y***_2_). GRFs that satisfy these two conditions are also referred to as second-order, wide-sense, or weakly homogeneous/stationary. Just as the name implies, a *homogeneous* GRF exhibits equivalent statistical properties at any given location. In addition, *F* is called *isotropic* if it is homogeneous and its covariance function only depends on the *distance* between its arguments (so that *C*(***x***_1_, ***y***_1_) = *C*(***x***_2_, ***y***_2_) whenever ||***x***_1_ −***y***_1_|| = ||***x***_2_ −***y***_2_||, where || · ||denotes geographic distance). Whereas a general homogeneous random field allows for the covariance function to depend on the spatial direction between its arguments, an isotropic random field absent of any such dependency on direction or orientation. When a GRF is isotropic, we write its mean and covariance functions respectively as *µ* and *C*(*d*), where *d* is the distance between two locations (e.g., *d* = ||***x*** −***y*** ||). Because *C*(0) is the variance of the random variable *F* (***x***) (for any ***x***), we call *V* = *C*(0) the *colocated variance* of the isotropic GRF *F*. The spatial *correlation* function is given by *ρ*(*d*) = *C*(*d*)*/V*. We therefore say the field *F* exhibits spatial autocorrelation whenever *ρ*(*d*)≠ 0 for some *d >* 0.

As an example of a GRF, consider a spatial white noise process *W*. Heuristically, we can think of *W* as an isotropic GRF with mean-zero, infinite colocated variance, and no spatial autocorrelation. Rigorously, setting *W*(*U*) = ∫_*U*_*W*(***x***)*d****x***, we have that *W*(*U*) and *W*(*V*) are normally distributed random variables with mean-zero, variances equal to the areas of the spatial regions *U* and *V* respectively, and covariance equal to the area of the intersection between *U* and *V*. Hence, if *U* and *V* do not overlap, *W*(*U*) and *W*(*V*) are independent random variables. The covariance function of a spatial white noise is the Dirac delta function *δ*(*d*) for which *δ*(*d*) = 0 and 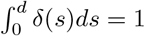 for all *d >* 0. In relation to the space-time white noise processes 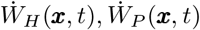 that appear in our model, we have 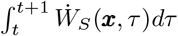 is a spatial white noise for both *S* = *H, P*.

An isotropic *k*-variate GRF ***F*** is composed of *k* isotropic univariate GRFs *F*_1_, …, *F*_*k*_. ***F*** is then completely characterized by means *µ*_1_, …, *µ*_*k*_ and covariance functions *C*_1_, …, *C*_*k*_ of the respective univariate fields *F*_1_, …, *F*_*k*_ along with the *cross*-covariance functions

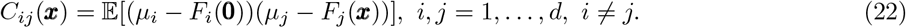

Similar to the colocated variance, we call *C*_*ij*_(0) the *colocated covariance* of *F*_*i*_ and *F*_*j*_ because it is the covariance of the random variables *F*_*i*_(***x***), *F*_*j*_(***x***) (for any location ***x***). Additionally, we call 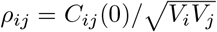 the *colocated correlation* of *F*_*i*_ and *F*_*j*_, where *V*_*i*_ = *C*_*i*_(0) is the colocated variance of *F*_*i*_.

### B Justification for Population Growth Rates

To begin, one may start with a pair of interacting individual-based branching processes where individuals are associated with a trait *z* ∈ ℝ and a geographic location *x* ∈ ℝ^2^. Assuming semelparous life-cycles, we model mortality and reproduction simultaneously so that individuals replace themselves with a Poisson number of offspring between unit intervals of time. The lifetime expected number of offspring (which, because mortality is equal for all individuals, we refer to as fitness) is determined by the trait of the parent along with the traits of other individuals the parent interacts with. This is similar to the starting points taken by Week et al. (2021) in the derivation of a diffuse-coevolution model and by Week and Nuismer (2021) in the derivation of the offset-matching coevolution model, except neither of those models have a spatial component.

To model fitness, we first consider the effects of abiotic selection 𝒜_*S*_ and biotic selection ℬ_*S*_ separately for host and parasite species (*S* = *H, P*). We decompose the effects of biotic selection into sources due to intraspecific competition 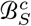 and interspecific parasitism 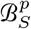 so that 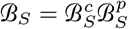. We will assume these effects multiply to produce the net fitness of an individual, *w*_*S =*_ 𝒜_*S*_ ℬ_*S*_. Set *w*_0, *S*_ the maximum fitness possible for species *S* in the absence of interspecific interactions and *θ*_*S*_ the abiotic optima trait value. Then, for either species, the multiplicative component of fitness due to abiotic stabilizing selection for an individual with trait *z* at any location is

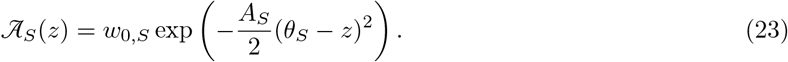

We assume competition occurs locally such that individuals that are geographically closer to each other induce stronger competition on one another than individuals that are farther apart. This induces a form of local population regulation that prevents run-away population growth. Denoting the distance between two spatial positions ***x*** and ***y*** by 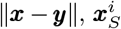 the location of the *i*th individual in species *S*, and *n*_*S*_ the number of individuals in species *S*, we model the effect of intraspecific competition on the *j*th individual as

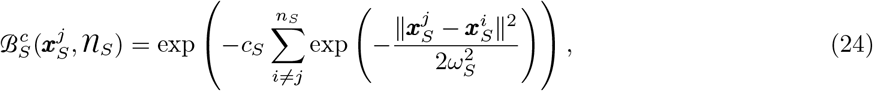

where *c*_*S*_ denotes the strength of spatial competition, *ω*_*S*_ is the spatial scale of competition, and 𝓃_*S*_ denotes the abundance measure for species *S*. Because 𝓃_*S*_(*U, V*) returns the number of individuals in species *S* with trait values in *U* ⊂ ℝ spatially located in the region *V* ⊂ ℝ^2^, 𝓃_*S*_ captures the spatial locations and trait values of individuals and hence the trait distribution and abundance in any region of space for species *S*.

We model host-parasite interactions by assuming a probability of infection that is a function of trait values given an encounter has occurred. Assuming a host individual with trait *z*_*H*_ encounters a parasite with trait *z*_*P*_, the probability of infection *α*(*z*_*H*_, *z*_*P*_) can be written as

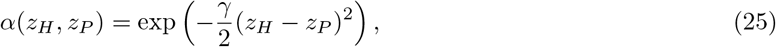

where *γ* ≥ 0 determines the sensitivity of this probability to differences in individual trait values. We will always assume weak sensitivity (ie, *γ* ≪ 1) so that *α*(*z*_*H*_, *z*_*P*_) ≈ 1− *γ*(*z* _*H*_−*z*_*P*_)^2^*/*2. We model the probability of encounter *ε* similarly as a function of the geographical distance between individuals. Denoting *ι* ≥ 0 the geographic scale of host-parasite interactions, we model the probability of encounter as

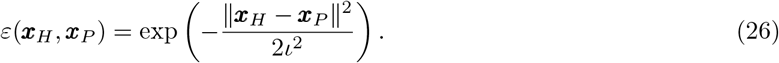

We allow *ι* ≪ 1 so that encounters may strongly depend on distance. Set *E*_*ij*_ the Bernoulli random variable representing whether the *i*th parasite encounters the *j*th host and *I*_*ij*_ the Bernoulli random variable representing the *i*th parasite infecting the *j*th host given their encounter. Assuming the parasite acquires the benefit *s*_*P*_ ≥ 0 and the host receives the cost *s*_*H*_ ≥ 0, the multiplicative effects of this single interaction on the fitnesses of the respective participants are exp(*s*_*P*_ *E*_*ij*_*I*_*ij*_) and exp(−*s*_*H*_*E*_*ij*_*I*_*ij*_). Taking expectations provide

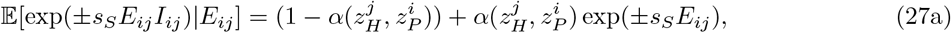

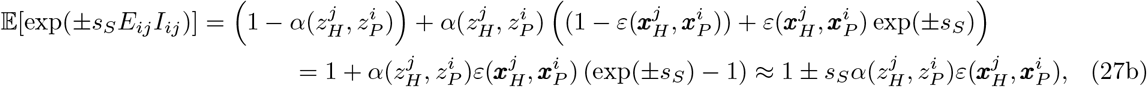

where the approximation holds when *s*_*H*_, *s*_*P*_ ≪ 1, which we assume from hereon. Then, assuming every parasite can potentially infect every host, the components of biotic selection due to interspecific interactions for each species are approximated by

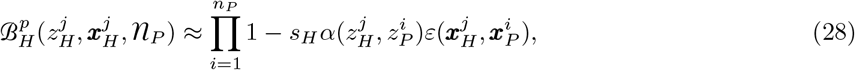

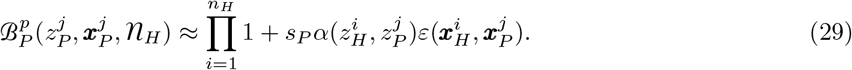

Using our weak biotic selection assumption *s*_*H*_, *s*_*P*_ ≪ 1, it will be convenient to rewrite these expressions as

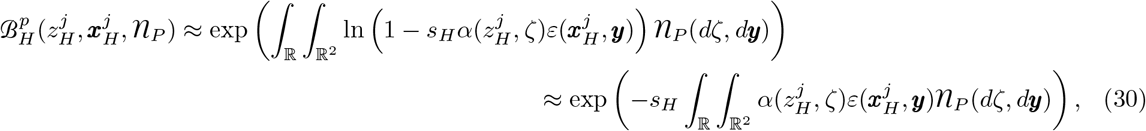

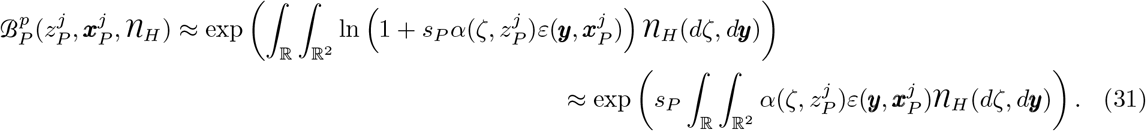

To model mutation and spatial movement, we assume offspring trait values are normally distributed around their parental value (technically, this is done with breeding values, see Week et al. 2021) and offspring locations are bivariate normal around their parental locations with i.i.d. displacements in the two spatial dimensions.

To take a diffusion limit of this individual-based process, we follow Week et al. (2021). In particular, for the *k*th stage of rescaling, the time interval between generations is divided by *k* (so it goes to zero as *k*→ ∞), the number of initial individuals *n*_*S*_ (0) in each species *S* = *H, P* is multiplied by *k* (so *n*_*S*_ (0) → ∞ as *k* → ∞), the variances of mutation and dispersal are divided by *k* (so both go to zero as *k*→ ∞), fitness for each individual is taken to the 1*/k*th power (so individual fitness tends towards unity as *k*→ ∞), and the *mass* of each individual is divided by *k* (so initial population *mass* remains *n*_S_ (0) for all *k* ≥ 1). In particular, this last part of our rescaling implies

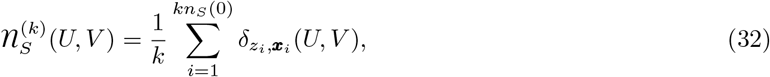

where *z*_*i*_ is the trait value of the *i*th individual, ***x***_*i*_ is its geographic location, and 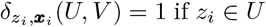 and ***x***_*i*_ ∈ *V* and zero otherwise. Sufficient conditions under which the rescaled individual-based process 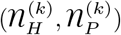 converges to a population-level process (𝔑_*H*_, 𝔑_*P*_) as *k* → ∞ are provided by Theorem 1 of Méléard and Roelly (1993). In particular, their condition (ℋ_1_) requires the sequence 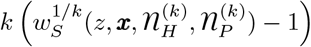 to converge to the population growth rate *m*_*S*_(*z, x*, 𝔑_*H*_, 𝔑_*P*_) of the population-level process 𝔑_*S*_ for species *S*. That is, population growth rates in the diffusion-limit are given by

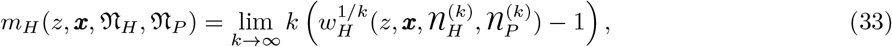

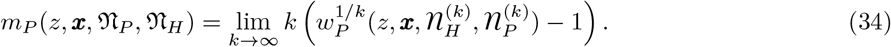

For the host we have

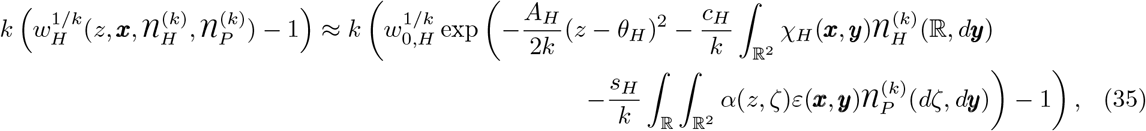

where we have set 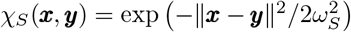. For large *k*, this is approximated by

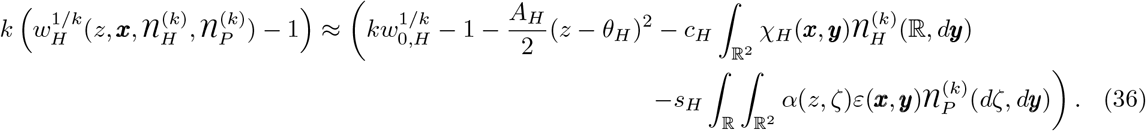

Then, setting *r*_*S*_ = ln *w*_0,*S*_ (the intrinsic growth rate for species *S*), we get

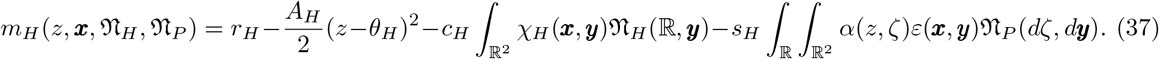

A similar expression for the parasite is also obtained. We now make the approximation that competition and selection are sufficiently weak relative to the intrinsic growth rate (i.e., *c, A*, ≪ *s r*) so that spatial fluctuations in local abundance densities due to selection are small relative to average local abundance density for each species when the system has reached stationarity. This implies the population growth rates *m*_*H*_, *m*_*P*_ are near zero when the system has reached stationarity. With this approximation, we write *N*_*S*_ as the abundance density for species *S* so that 𝔑_*S*_(ℝ, *U*) ≈ |*U* | *N*_*S*_, where ℝ is given as the first argument to N to include individuals with any trait value and |*U* | is the area of the geographic region *U* ⊂ ℝ^2^. In this case we have

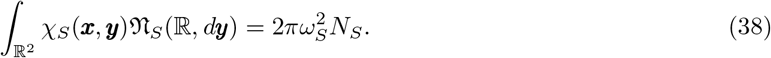

The term capturing the effects of the host-parasite interaction has an integral across phenotypic space and an integral across geographic space. To simplify the geographic integral, set

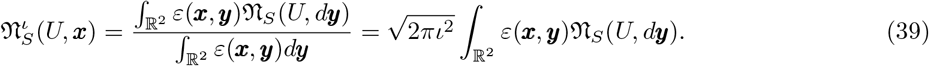

This notation makes sense because *ε* is a smooth integrable function and a convolution with such a function yields another smooth function. Furthermore, when 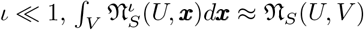. Using our assumption that *γ* ≪ 1, the biotic and abiotic components cumulatively contribute quadratic selection. Given that stabilizing abiotic selection is sufficiently strong relative to disruptive biotic selection on the host, trait distributions at any location will be approximately normal with mean and variance 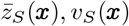 for species *S* at location ***x***. Then, assuming *ι* ≪ 1, this implies

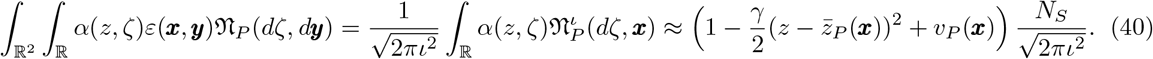

Because selection is quadratic and abundance is constant, selection and drift decay phenotypic variance at a constant rate. From our assumption of Gaussian mutations, phenotypic variance also has a constant rate of input. We can therefore expect phenotypic variance for each species to eventually fluctuate stochastically around a spatially constant equilibrium. We thus further approximate by setting the phenotypic variances equal to those constant equilibria. We can therefore approximate the growth rates for each species as

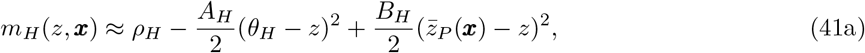

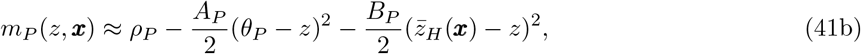

where we have dropped the dependencies on 𝔑_*H*_, 𝔑_*P*_ for brevity and set 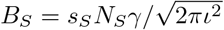 and

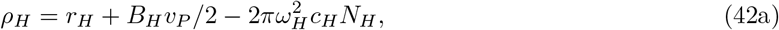

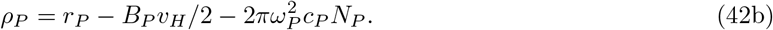

### C Gaussian Distribution of Local Traits

To show that local trait distributions can be approximated by Gaussian distributions, we start by considering deterministic dynamics of single species experiencing abiotic stabilizing selection. This leads to a deterministic partial differential equation describing the dynamics of the trait distribution and involves diffusion in both trait space and geographic space. The equilibrium solution is spatially homogeneous, allowing us to focus on characterizing the trait distribution at a single location. After confirming the equilibrium trait distribution is Gaussian at all locations for a single species, we move on to incorporate interspecific interactions following our model of host-parasite trait-matching. This leads to a pair of interacting partial differential equations generalizing the single species equation mentioned above. Again, the equilibrium solution is spatially homogeneous, so we focus on a single location. After confirming the trait distribution is Gaussian for interacting species case, we then argue that for sufficiently large population sizes in which the diffusion approximation holds, random genetic drift should only slightly perturb the trait distribution. Hence, local trait distributions should be approximately Gaussian and, therefore, approximately free of skew.

We begin the dynamics of the abundance density *ν*(*z*, ***x***) (where *N* (***x***) = ∫_ℝ_*ν*(*z*, ***x***)*dz* is the total abundance at location ***x***). We assume Gaussian descendants so that offspring traits are normally distributed around parental traits with variance *ε*^2^. This provides

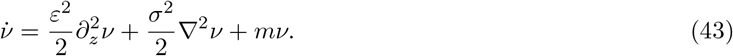

Following Appendix B, the growth rate for a single species under our model that is not engaged in an interspecific interaction is given by

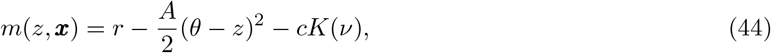

where

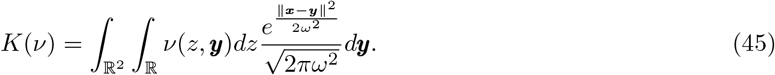

One can check that 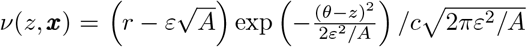 satisfies equation (43). In particular, this implies the equilibrium trait distribution is spatially homogeneous and Gaussian with mean 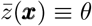 and variance *v*(***x***) ≡ *ε*^2^*/A* along with spatially homogeneous local abundance 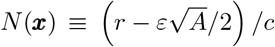. Furthermore, at sufficiently large local abundances for which the diffusion approximation outlined in Appendix B is valid, demographic stochasticity and random genetic drift will have only small effects on the trait distribution relative to the effects of diffusive mutation and quadratic stabilizing selection so that the trait distribution remains approximately Gaussian.

For the case of host-parasite interaction mediated by a trait-matching model, we can rewrite the growth rates as

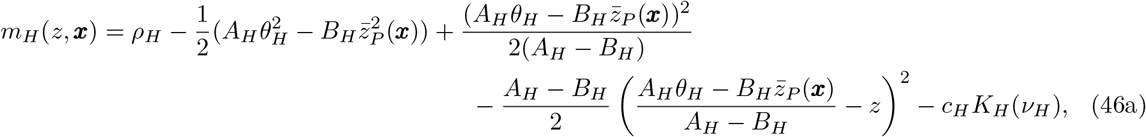

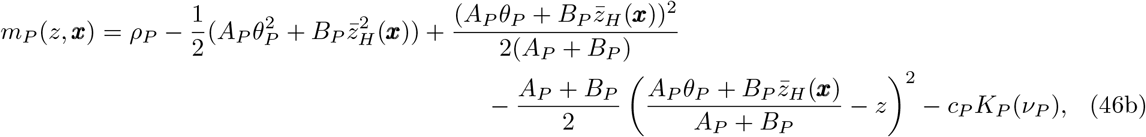

with *ρ*_*H*_, *ρ*_*P*_ given by equations (42). Although these growth rates appear more complicated, they are still quadratic functions of the focal species’ trait value. Hence, equilibrium trait distributions will be spatially homogeneous and Gaussian for both species. In particular, these equilibrium trait distributions are free of skew that appears when geographic structure is discrete and abiotic optima are spatially variable (Débarre et al. 2015). In the stochastic case, our assumptions of weak selection and large local abundance prevent significant feedbacks between evolutionary and abundance dynamics and imply phenotypic variation occurs at sufficiently large spatial scales relative to dispersal so that local trait distributions should remain approximately normal.

